# Towards improved particle averaging for single-molecule localization microscopy using geometric deep learning

**DOI:** 10.1101/2025.11.26.690737

**Authors:** Diana Mindroc-Filimon, Dominic Helmerich, Patrick Salome, Markus Sauer, Philip Kollmannsberger

## Abstract

Single-molecule localization microscopy (SMLM) can resolve intracellular structures down to the nanoscale, but often produces sparse and incomplete data. Particle averaging (PA) can aid with the reconstruction of complete structures, but traditional PA methods can suffer from template bias or the high computational costs of geometric alignment. To address these limitations, we developed a geometric deep learning (GDL) framework for enhanced, template-free 3D particle averaging. Our pipeline uses a GDL autoencoder, trained on realistic simulated data, to map incomplete point clouds into a robust latent space. By averaging feature vectors directly within this space, our method bypasses the need for explicit 3D alignment. We validated our approach on simulated DNA origami and experimental nuclear pore complex (NPC) data. The latent space averaging successfully reconstructed NPC structures with key metrics (ring radius ≈ 46 nm, ring distance ≈ 52 nm) that are comparable to state-of-the-art methods. This work establishes a viable GDL pipeline for SMLM analysis, offering an efficient alternative to traditional PA. While the current model requires structure-specific training, our results highlight the significant potential of GDL for quantitative structural biology.

## Introduction

The emergence of super-resolution microscopy, in particular single-molecule localization microscopy (SMLM), has reshaped cell biology by enabling the visualization of sub-cellular structures at nanoscale resolution (1, 2). SMLM techniques can resolve individual molecules with precisions down to 20 nm. This capability provides unprecedented opportunities for the quantitative analysis of protein distributions and the structural characterization of molecular complexes *in situ*, with demonstrated applications in neurodegenerative disease research and oncology (3, 4).

However, these super-resolution methods are fundamentally limited by the inherent incompleteness of the acquired data. The stochastic nature of fluorophore activation and detection, combined with challenges in achieving dense and uniform labeling, often results in sparse and fragmented representations of the underlying biological structures (5, 6). This undersampling hinders the complete reconstruction of molecular architectures and complicates quantitative analysis.

To overcome this challenge, particle averaging (PA) has been widely adopted as a computational strategy to enhance the signal-to-noise ratio and structural completeness (7). Originating from cryo-electron microscopy (8), PA combines multiple observations of the same particle, which are often noisy or incomplete, to generate a high-fidelity average. However, conventional PA methods for SMLM data have significant drawbacks. Many PA alogrithms are image-based, requiring the conversion of 3D point cloud data—the direct output of SMLM methods—into a gridded format. This transformation can introduce discretization artifacts and a loss of precision (9). Furthermore, template-based alignment methods risk biasing the final reconstruction towards the chosen template, potentially obscuring novel or heterogeneous structural features (10).

The intrinsic format of SMLM data as 3D point clouds motivates a shift towards computational tools designed for such non-Euclidean data. Geometric deep learning has recently emerged as a powerful paradigm for analyzing and interpreting complex geometric structures (11). Architectures like PointNet (12) and its successors can operate directly on point clouds, respecting their permutation invariance and spatial nature. This opens a promising avenue for developing template-free and alignment-agnostic particle averaging workflows. However, the application of geometric deep learning to SMLM data is still incipient and faces challenges, including the lack of ground-truth data to train supervised models and the need to adapt existing architectures to the sparse and noisy nature of biological datasets (13).

In this work, we propose a novel framework that leverages geometric deep learning to enhance 3D particle averaging for SMLM, with the aim of reconstructing complete structures from underlabeled or partial observations. Our approach circumvents the need for explicit 3D alignment by encoding particles into a rotation-invariant latent space. We hypothesize that these learned representations can be used to effectively classify heterogeneous structures and that averaging them can robustly reconstruct a complete particle. To enable model training, we first developed a simulation framework to generate realistic SMLM datasets. Our contributions are twofold: we 1) present a method for learning compact and descriptive latent representations directly from the sparse point clouds characteristic of SMLM data, enabling accurate structural classification; and we 2) introduce a particle averaging pipeline that operates within this latent space, thereby eliminating template bias and the computationally intensive alignment step. We demonstrate that this approach can faithfully reconstruct complete biological structures from multiple incomplete instances, offering a significant advancement over state-of-the-art methods. A graphical abstract of the workflow is presented in Fig. 1.

**Fig. 1.**
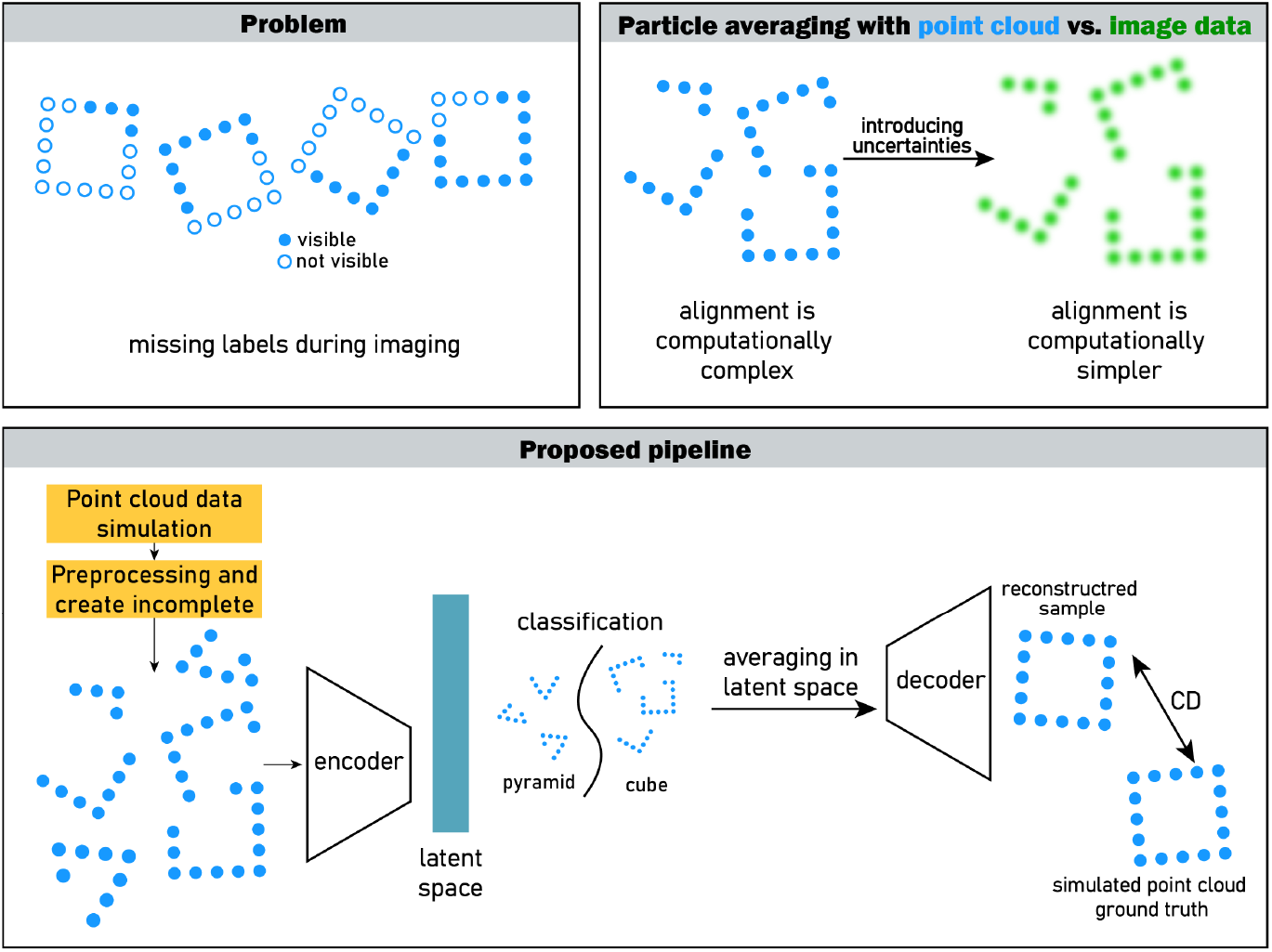
Proposed pipeline. Transforming point cloud data into image data introduces additional uncertainties, motivating our choice of working directly with point cloud data. Typically, alignment of point cloud data is more computationally complex than the alignment of image data. Therefore we decided to work with permutation invariant networks. After data simulation, preprocessing and creating incomplete samples, the point cloud deep learning network is trained for reconstruction and classification in latent space simultaneously. Multiple encoded incomplete samples are averaged and then decoded for obtaining a complete, higher-resolution structure. CD: Chamfer Distance. Encoder and decoder components are detailed in later sections.

### Background and related work

Many particle averaging approaches are template-based, aligning either rendered images (9, 14) or coordinate data (15) to a predefined structural model, which risks introducing bias. To address this, state-of-the-art methods have shifted towards template-free, coordinate-based particle fusion (7). These techniques perform iterative registration of multiple point clouds, often using Gaussian Mixture Models, to determine particle poses before averaging (10, 16). While powerful, these registration-centric methods can be computationally intensive and sensitive to initialization. In parallel, deep learning has been applied to SMLM, but predominantly for image-based tasks like emitter localization from raw camera frames (17), not for particle averaging of reconstructed point clouds. The direct application of geometric deep learning to SMLM point clouds for structural analysis remains an emerging field. While pioneering work has used PointNet-based architectures for the specific sub-task of particle alignment (18), the potential of more advanced models to create an end-to-end particle averaging framework is largely unexplored. To our knowledge this work presents the only application of point cloud deep learning to SMLM particle averaging. Reconstructing a point cloud from its latent space representation presents unique challenges not found in image-based decoding, primarily due to the unstructured and variable-cardinality nature of point set data. A prominent architecture, FoldingNet (19), addresses this by generating a 3D point cloud by learning a continuous mapping that “folds” a canonical 2D grid into the target shape, guided by the latent vector. While successfully applied to model cell morphology (20), its efficacy is limited for structures with a non-planar topology. A more versatile approach, particularly for the complex assemblies found in SMLM, is the decoder from the Point Completion Network (PCN) (21). This architecture employs a two-stage, coarse-to-fine strategy: it first predicts a sparse point cloud representing the global structural skeleton and then generates dense, high-fidelity details by applying localized folding operations around these initial points. This hierarchical process is adept at capturing both the overall topology and the intricate local features of a structure.

The training of such decoders is guided by loss functions that measure the similarity between the generated and ground-truth point clouds. In the deep learning context, the dominant loss function is the Chamfer Distance (CD), which efficiently computes a symmetric, average distance between each point in one set and its nearest neighbor in the other. This optimization target stands in contrast to the metrics used in traditional, registration-centric particle averaging. In image-based alignment, the objective is typically to maximize a cross-correlation score in pixel space. In coordinate-based alignment, the goal is to minimize spatial distances, either to a fixed template using simple closest-point measures (9) or between pairs of particles. The latter often involves more sophisticated metrics that treat the particles as probability distributions, such as the Bhattacharyya distance between Gaussian Mixture Model representations, to achieve robust, template-free alignment (7).

## Materials and methods

### Experimental data

We used an experimental public dataset of macromolecular structures imaged with single-molecule localization microscopy, as described by Heydarian et al. (2021) (22). The dataset comprises two structures: DNA-origami tetrahedrons and nuclear pore complexes (NPCs). The DNA-origami dataset contains 498 tetrahedron-shaped nanostructures imaged using 3D astigmatic DNA-PAINT. The nanostructures were synthesized via a one-pot reaction with 10 nM scaffold strand (p8064), 100 nM each of core, connector, vertex, biotin, and DNA-PAINT handle staples, and 1400 nM biotin anti-handles. For imaging, 30,000 frames were acquired at an 800 ms integration time using a 2 nM P1-Cy3b imager, with an excitation intensity of 226 W/cm2 at 561 nm. The NPC dataset was generated by imaging the Nup107 protein labeled with Alexa Fluor 647 via a SNAP-tag. It includes two subsets, one acquired with 3D astigmatic DNA-PAINT containing 306 particles with an average of 88 localizations per particle and one with 4Pi-STORM containing 750 particles with an average of 58 localizations per particle.

### Training data

The training dataset includes 1000 samples of DNA origami tetrahedrons, each consisting of a set 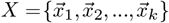 of *k* point coordinates, 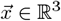. Leveraging statistical parameters extracted from an experimental dataset of 498 DNA-origami tetrahedra (22), we established probability distributions for key structural features, including the number of localizations per cluster and their standard deviations in three dimensions. This framework simulates predefined geometric scaffolds (e.g., tetrahedra) and populates them with point clouds, incorporating experimentally-derived statistics to enhance biological realism. To address reconstruction challenges, both anisotropic and isotropic point cloud samples were generated by sampling from fitted statistical distributions of the standard deviations. To ensure the simulation framework accurately replicated the statistical properties of real SMLM data, probability distributions were fitted to main experimental metrics extracted from the experimental dataset, specifically the number of localizations per structure, cluster sizes (localizations per binding site), and cluster standard deviations in x, y, and z. Several candidate distributions, including Lognormal, Gamma, and Weibull, were evaluated for goodness-of-fit using the Akaike Information Criterion (AIC). The fidelity of the simulated data to the experimental data was validated through comparative analysis using histogram overlays and statistical assessments, including the Kolmogorov-Smirnov and Mann-Whitney U tests, ensuring the synthetic data accurately reflects real-world experimental conditions.

A standardized data preparation pipeline was established to ensure consistency across diverse datasets for our deep learning applications. To enhance data quality, we performed denoising using HDBSCAN for its proficiency in handling variable density clusters and noise inherent in SMLM data. To simulate realistic biological scenarios of incomplete labeling, we implemented a strategy involving the random removal of entire point cloud clusters, identified by HDBSCAN, to achieve specific labeling efficiencies (e.g., 75% and 50%). Additionally, preprocessing included rotations and translations to increase data variability mimicking experimental conditions. We also implemented padding by duplication to standardize input dimensions, facilitating consistent batch processing while preserving data integrity. Figure 10 presents preprocessed data for training.

### Training loss

To evaluate the reconstruction quality, we use the widely established Chamfer Distance (CD) as a loss function to measure the distance between the reconstructed point cloud and the ground truth. The CD is a measure of the distance between two point sets *A* and *B*, calculated as the average distance between points in *A* to the closest point in *B* and vice versa:

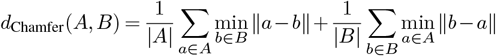

### Training details

The model was implemented using PyTorch and trained on an Nvidia Quadro RTX A5000 GPU with 24GB of memory. We used a 70/20/10 split to divide our dataset into training, validation, and test sets. The model was trained using the Adam optimizer with an initial learning rate of 0.001 and a batch size of 64. The momentum was set to 0.9, and momentum2 to 0.999. The training was carried out for a maximum of 500 epochs, with an early stopping patience of 30 epochs. A ReduceLROnPlateau scheduler was employed, which reduced the learning rate by a factor of 0.75 if the validation loss did not improve for 10 consecutive epochs. All code for data simulation, network training and validation is freely available under https://github.com/dianamindroc/smlm.

## Results

### Data simulation

The dataset used in this work consists of simulated DNA-origami data. DNA origami involves the folding of DNA to create 2D and 3D objects at the nanoscale (23). Its predictable and programmable nature en-sures that the resulting structures are consistent and reproducible, thus they provide an excellent source of ground truth data for the development and evaluation of computational models. A custom data simulation framework was developed to generate realistic synthetic datasets for model training and evaluation. The Lognormal distribution consistently yielded the lowest AIC values across all metrics, identifying it as the optimal model for generating realistic synthetic data. The Kolmogorov-Smirnov (KS) test confirmed no statistically significant difference between the datasets (*p >* 0.05), thereby verifying the simulation’s suitability for supervised network training.

Samples of simulated tetrahedron are presented in Fig. 2, displaying structural fidelity to the experimental data in (10).

**Fig. 2.**
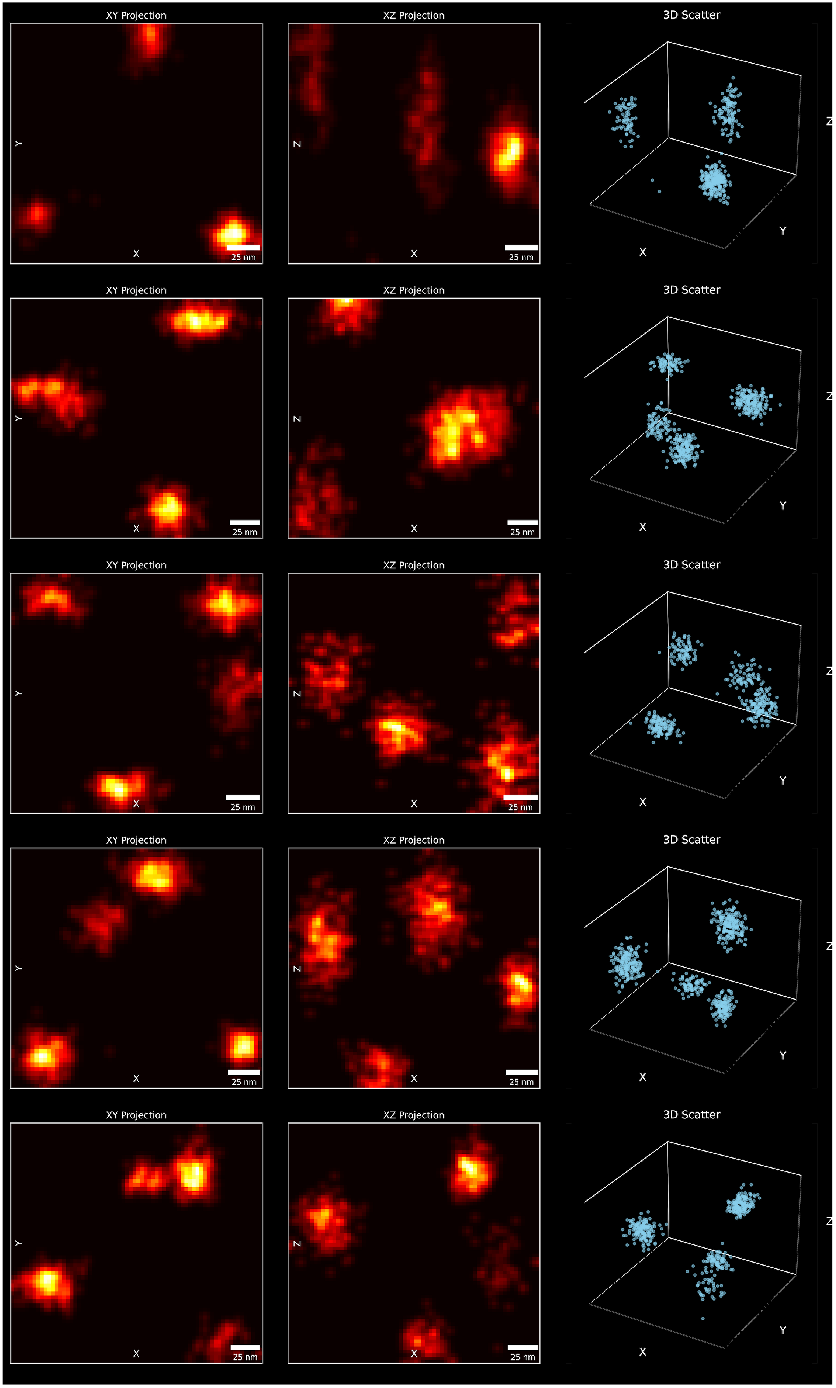
Simulated DNA-origami tetrahedron samples; Left: 2D projections of the simulated tetrahedron point clouds onto the XY and XZ planes, right: 3D scatter plot of the same tetrahedron sample, providing a volumetric view of the localization clusters

### Model selection

To identify the optimal encoding architecture for SMLM data, a comparative analysis of five leading neural network models was performed. We evaluated AtlasNet (24), FoldingNet (19), Point Completion Network (PCN) (21), PointNetFCAE (25), and TopNet (26) based on reconstruction fidelity, computational performance, and pose-invariance. Each model was initially trained on the large-scale ShapeNet benchmark dataset as intended by their original design. Subsequently, the pre-trained models were tasked with completing partial structures from our experimental DNA-origami dataset. The primary evaluation criterion was reconstruction accuracy, which was quantified using the Chamfer Distance between the generated output and the ground truth point cloud.

The results of the comparison study for choosing the point cloud deep learning architecture that works best on SMLM data are reported as CD across the test set and presented in Table 1.

**Table 1.** Results of the comparison study of different state-of-the-art point cloud deep learning architectures on DNA origami. The reported measure is the Chamfer Distance between reconstruction and ground truth for the point completion task with 95% confidence interval based on t-distribution.

| Architecture | Chamfer Distance |
| --- | --- |
| FoldingNet | 1.98 [1.79 - 2.11] |
| **PCN** | **0.16 [0.12 - 0.18]** |
| TopNet | 8.37 [7.98 - 8.88] |
| AtlasNet | 7.85 [7.21 - 8.03] |
| PointNetFCAE | 7.99 [7.42 - 8.23] |

From the models evaluated, the Point Completion Network (PCN) was selected as starting point for further refinement.

### Point cloud deep learning for SMLM

We developed Pocafoldas, a network tailored for the reconstruction and analysis of SMLM point cloud data. The network’s encoder is founded on a hierarchical feature learning strategy inspired by PCN (21). It first processes the input point cloud with a convolutional network to extract point-wise features, which are then aggregated via max-pooling to form a global feature descriptor. This global descriptor is concatenated back with the initial point-wise features to create a rich hybrid representation that preserves both local geometric detail and global context. This representation is processed by a self-attention mechanism, which adaptively refines the feature space by emphasizing structurally important points. The final 1024dimensional latent vector is generated using mean-pooling, a deliberate choice over max-pooling to create a global descriptor that better reflects the overall distribution and density of the localizations.

The decoder implements a two-stage, coarse-to-fine reconstruction process. It first uses a multi-layer perceptron (MLP) to transform the global feature vector into a coarse structural scaffold. This scaffold is then refined using an adaptive folding mechanism, where a canonical 2D grid is deformed into a detailed 3D structure based on the learned global features. We modified the decoder to dynamically adjust the size of the output point cloud to match that of the input. Unlike standard models with fixed outputs, it preserves the quantitative nature of the localizations and prevents the network from hallucinating points not present in the experimental data. The architecture and data flow pipeline are visualized in Fig. 3.

**Fig. 3.**
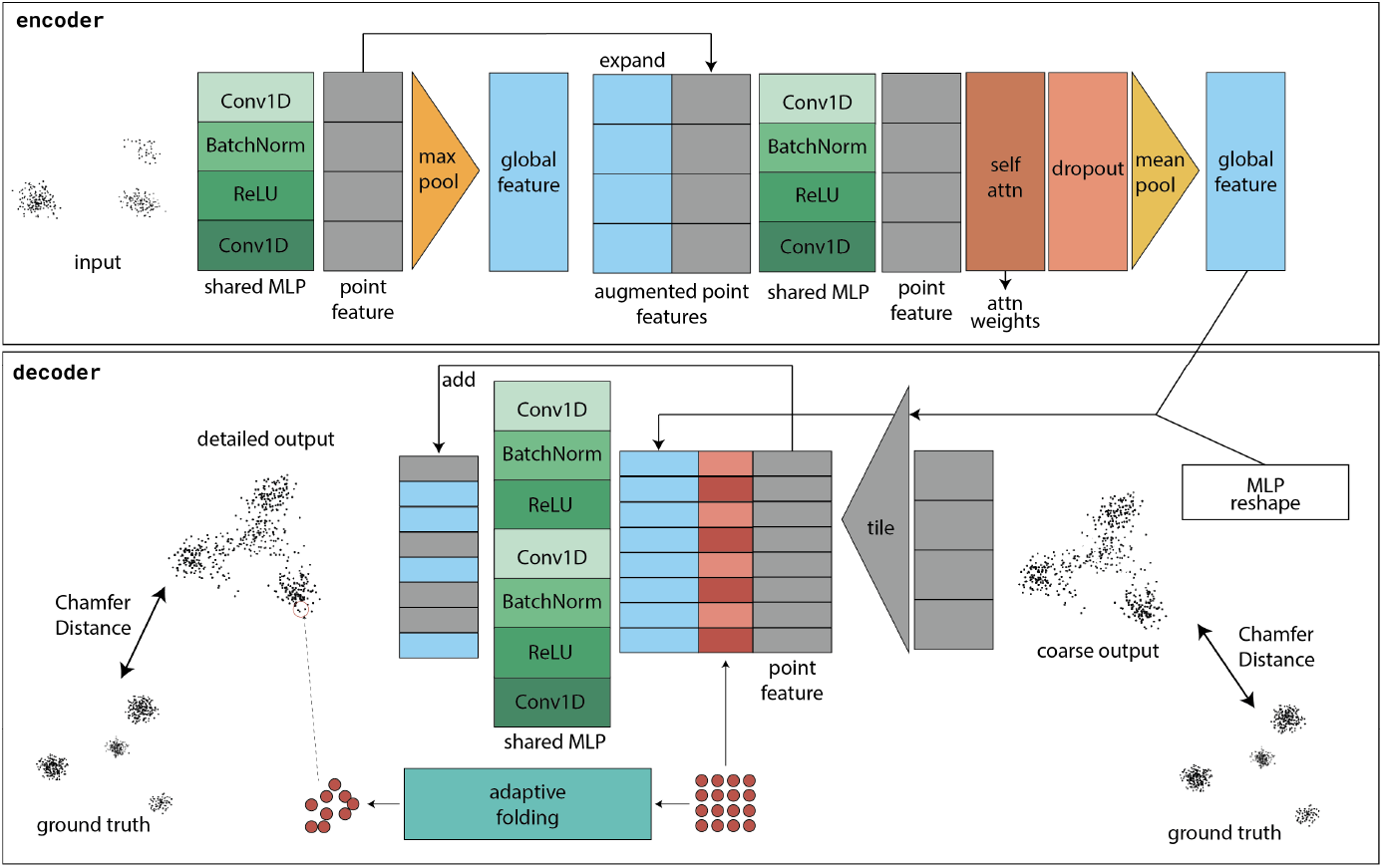
Architecture of the Pocafoldas network. The model processes input point clouds through a series of shared MLPs and pooling operations to extract global features. We introduced key improvements including dropout and self-attention mechanisms to enhance feature learning and model generalization. The network produces both coarse and detailed outputs, with the latter refined through an improved folding operation, called adaptive folding. This operation allows for more accurate reconstruction of complex geometries. Both outputs are compared against ground truth data using Chamfer distance as a metric.

### Clustering and classification in latent space

To investigate the quality of the learned representations, we visualized the 1024-dimensional latent space of encoded simulated DNA-origami cubes and tetrahedra using t-Distributed Stochastic Neighbor Embedding (t-SNE) (27). To test the robustness of the encoding against incomplete data, we generated t-SNE projections for structures with 0, 1, and 2 clusters systematically removed.

Our encoding strategy successfully generates a robust latent representation that distinguishes between different geometries while remaining invariant to missing data. As illustrated in the t-SNE projection (Fig. 4), the encoded cube and tetrahedron structures occupy distinct, well-separated regions of the latent space. This separation persists even after the removal of one or two clusters - within each class, the representations of complete and incomplete structures cluster together, confirming that the model effectively recognizes them as variants of the same object.

**Fig. 4.**
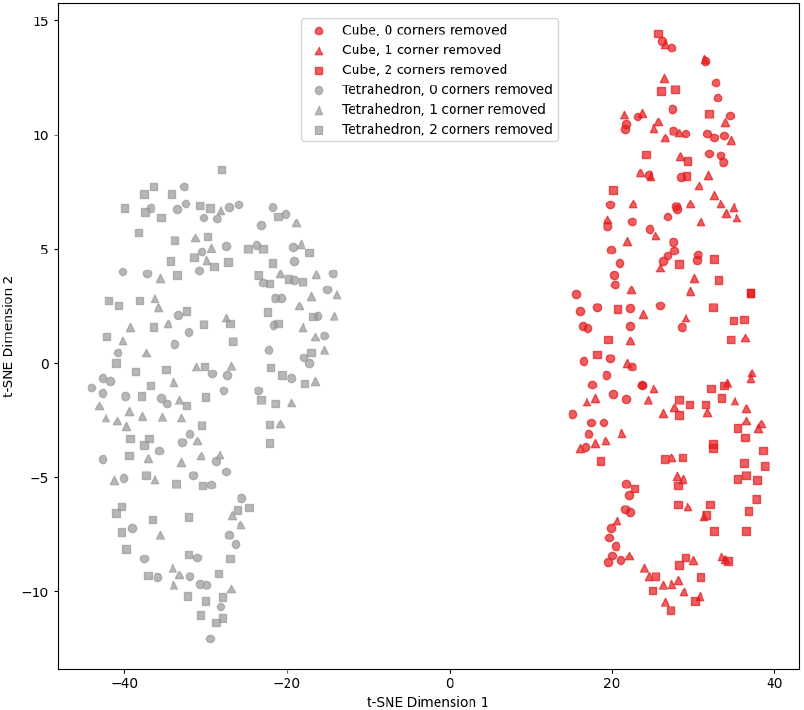
t-SNE visualization of DNA origami structures. The plot shows the distribution of cube (red) and tetrahedron (gray) structures in a two-dimensional space derived from their high-dimensional feature representations. Each point represents a single structure, with different shapes indicating the number of corners removed: circles (0), triangles (1), and squares (2). The clear separation between red and gray clusters demonstrates that the model effectively distinguishes between cubes and tetrahedron. Within each structural class, the overlapping distribution of different shapes suggests that the model is robust to corner removal, treating structures with varying numbers of removed corners as similar.

### Systematic experimentation

We conducted a systematic evaluation of the network architecture through two distinct training approaches: autoencoding and point cloud completion.

The initial autoencoder training was designed to assess the model’s ability to create high-fidelity reconstructions of its own input and to generate a meaningful latent space. In this phase, we challenged the network with various data types, including simulated point clouds with both isotropic and anisotropic distributions, multiple sample sizes, and distinct geometries (tetrahedra and cubes).

In the subsequent point cloud completion training, the network was tasked with a more complex objective: reconstructing complete, isotropic ground truth structures from incomplete and anisotropic inputs. To identify the most effective architecture, we performed a systematic, incremental evaluation of key components. We began with a baseline model and sequentially integrated an adaptive folding mechanism, a self-attention module, and a dropout layer. An ablation study was also conducted by removing the folding mechanism entirely to assess the network’s capacity for direct point generation. The optimal model configuration was first determined through rigorous training on our simulated DNA-origami dataset. We then applied the adapted network on simulated incomplete underlabelled DNA origami data with anisotropic noise. An example of the DNA origami tetrahedron from the reconstruction results on the test set with the best performance can be seen in Fig. 5, with CDs of 0.2521, 0.2375 and 0.2267 respectively. This best-performing architecture was then subsequently trained and evaluated on the experimental NPC dataset. Completion results are presented in Fig. 6.

**Fig. 5.**
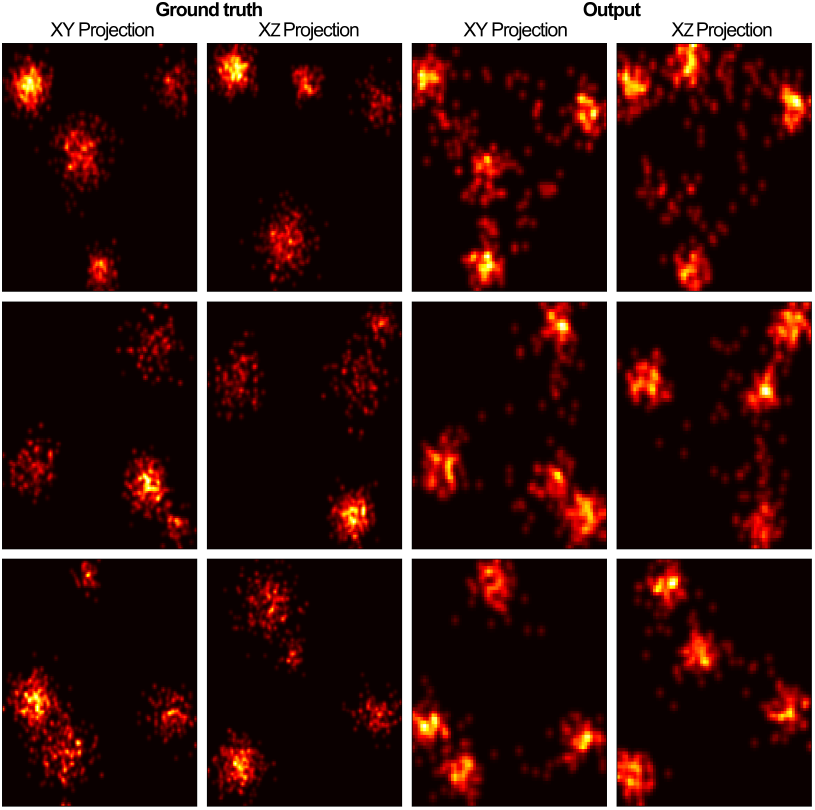
DNA origami completion results. Left: Ground truth sample, XY and XZ projections. Right: Reconstructed sample, XY and XZ projections. The reconstructed outputs show high similarity to the ground truth, accurately restoring the 3D positions and morphology of the DNA-origami assemblies, highlighting the model’s ability to perform high-fidelity structural completion.

**Fig. 6.**
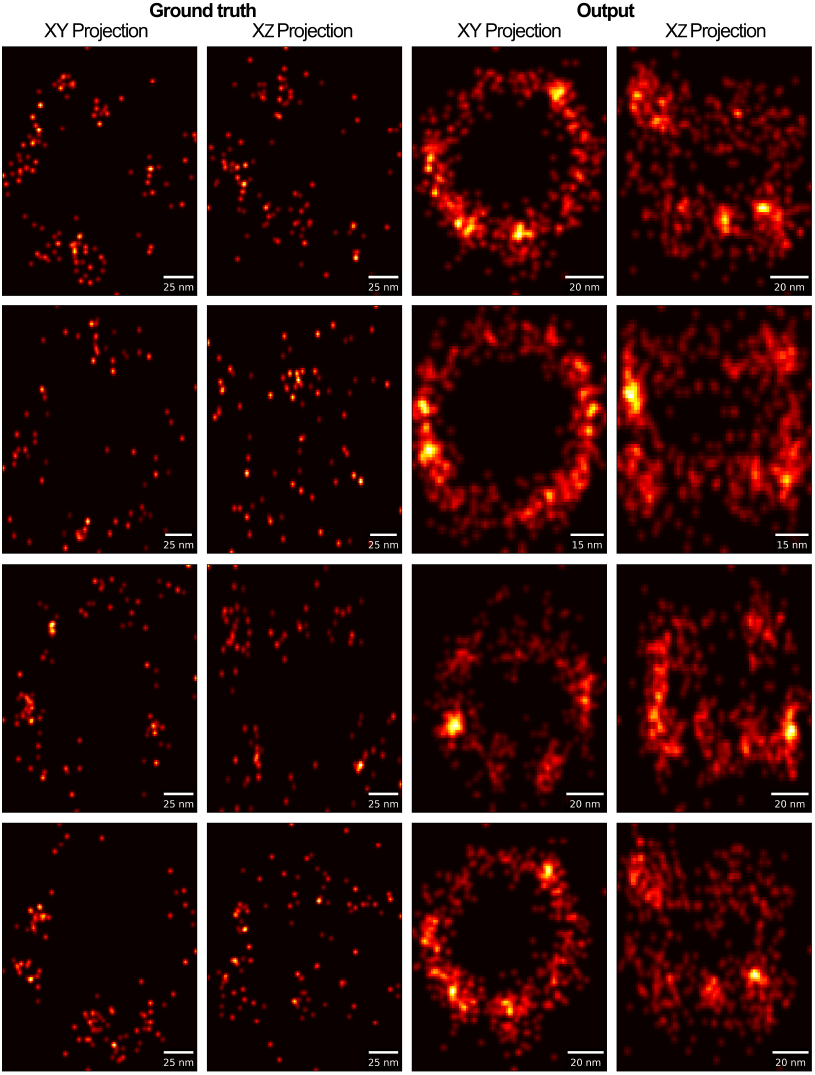
Nuclear pore complex reconstruction results. Left: input point clouds, XY and XZ projections. Right: reconstructed point clouds, XY and XZ projections

### Particle averaging

Following encoding, the SMLM point clouds are averaged in latent space. This process leverages the permutation-invariant nature of the encoded representations, circumventing the challenge of establishing point-to-point correspondences required by traditional averaging methods. Rather than decoding the direct mean of the latent vectors, we identify a medoid: the single feature vector from the original dataset closest to the average representation, as determined by the squared Euclidean distance. This medoid is then decoded to generate the final, averaged structure, ensuring that the resulting point cloud is a high-likelihood representation from the learned data manifold, which mitigates the risk of generating artifacts from decoding a vector that lies outside the distribution of observed structures.

As illustrated in Fig. 7, the decoded medoid successfully reconstructs the canonical architecture of the NPC. The model resolves the distinct cytoplasmic and nucleoplasmic rings, forming the well-known double-ring structure. Furthermore, the observed ring spacing and approximate symmetry are consistent with known NPC morphology, underscoring the structural accuracy of the averaging process.

**Fig. 7.**
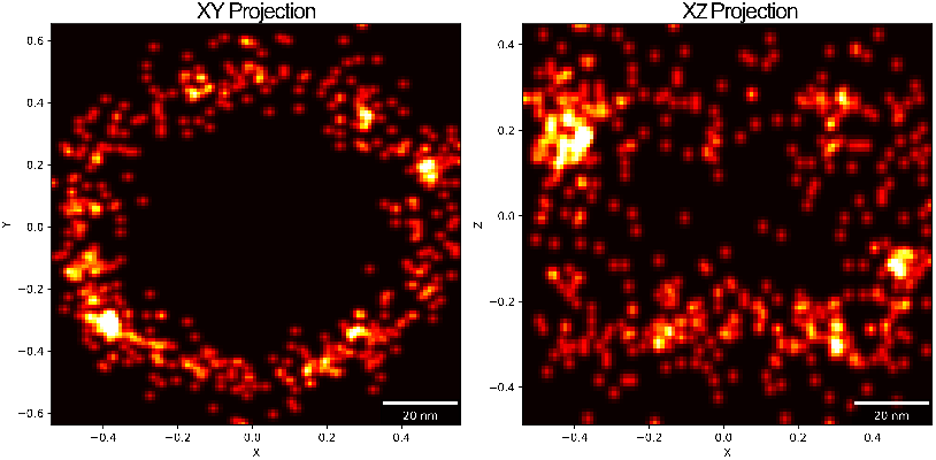
**Decoded NPC closest to the average feature vector** obtained by averaging all encoded feature vectors from the test set. Scale bar: 20nm

### Quantitative comparison

The performance of the methods developed in this work was benchmarked by comparing key structural metrics against template-free all-to-all registration as described in Heydarian et al. (10) and cryo-EM measurements. These metrics were extracted from the average particle structure (i.e., the decoded medoid, Fig. 7). The analysis included characterizing ring diameters from the radial distribution of localized points (Fig. 8) and determining the position and separation of stacked rings from their axial distribution, as represented by the z-coordinate histogram (Fig. 9).

**Fig. 8.**
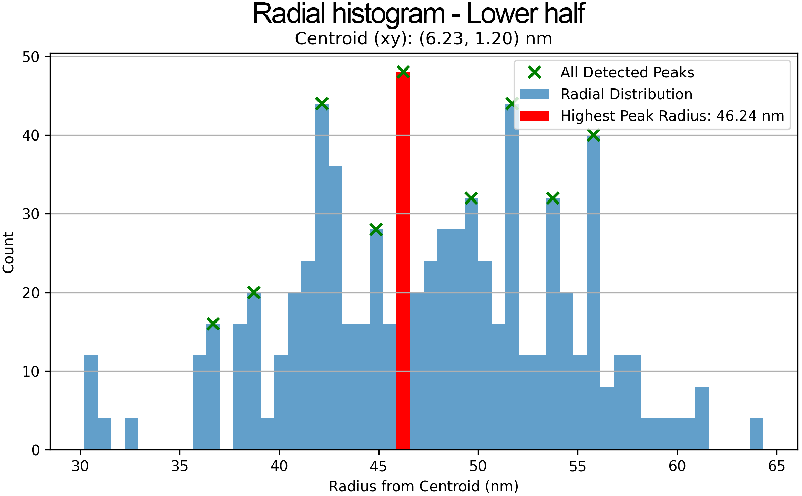
Radial distribution analysis for the nuclear ring of the reconstructed NPC. The histogram shows the count of points as a function of radius from the centroid (6.23, 1.20 nm). Detected peaks are marked with green ‘x’, with the highest peak (red bar) identified at a radius of 46.24 nm.

**Fig. 9.**
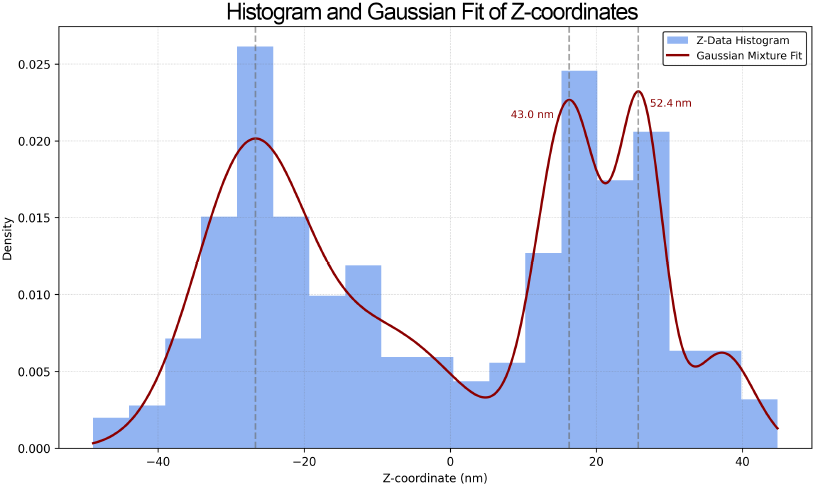
Z-histogram of reconstructed NPC. A Gaussian Mixture Model fitted to the distribution reveals a multi-modal profile characteristic of the NPC’s architecture. The distance between the primary peaks corresponds to a ring separation of 52.4 nm, consistent with the expected spacing between the cytoplasmic and nuclear rings. An intermediate peak (at 43.0 nm separation) is attributed to a local density hotspot within the reconstructed point cloud, likely an artifact.

The radial histogram indicates a nuclear pore complex (NPC) radius of 46 nm (Fig. 8). The z-histogram reveals a distance of 52.4 nm between the cytoplasmic and nuclear rings (Fig. 9).

In Table 2, we compare the metrics derived with our method to those reported by Heydarian et al. (10), who applied a template-free averaging method, and to data from the high-resolution cryo-EM structural map of the human NPC scaffold by von Appen et al. (28).

**Table 2.**
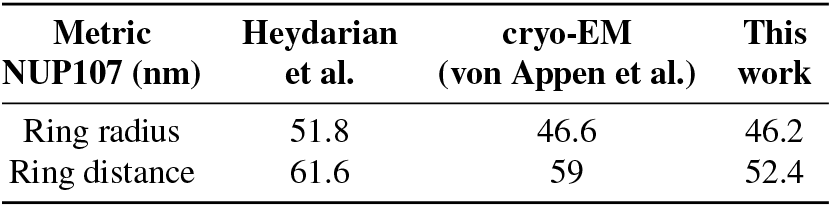
Comparison of structural metrics for NUP107. Ring radius is the radius of the nuclear ring and ring distance is the distance between cytoplasmic and nuclear rings.

## Discussion

This work introduces a novel geometric deep learning pipeline to address main challenges in 3D single-molecule localization microscopy particle averaging, namely data incompleteness and the computational cost of traditional alignment. Our innovation is a method that performs particle averaging directly in a learned latent space, bypassing the need for explicit point cloud registration. This approach provides a computationally efficient and robust framework for reconstructing complete structures from sparse, noisy observations.

The main element of our pipeline is the Pocafoldas network, an encoder-decoder architecture specifically tailored for SMLM data. To enable supervised training, we first developed a custom simulation framework that generates realistic point clouds by sampling from statistical distributions derived from experimental DNA origami data. The lack of ground truth information in experimental localization microscopy data makes it challenging to train accurate models. Therefore, using simulations can provide a controlled environment to generate high-quality training data, hence the approach provided annotated datasets with corresponding ground truth. The Pocafoldas architecture was then systematically optimized through key modifications: a self-attention mechanism to focus on geometrically salient features, mean pooling to generate a more holistic global feature descriptor, and a folding-based decoder with dynamic output sizing to reconstruct the structure with a natural point density. These adaptations were aimed at learning meaningful structural representations from the sparse and incomplete point clouds characteristic of SMLM.

Our primary contribution is particle averaging that operates within the permutation-invariant latent space learned by Pocafoldas. Instead of performing iterative geometric alignment on raw coordinates, our method averages the latent space vectors of all particles and identifies the medoid (the latent vector of an observed particle closest to the average). Decoding this medoid vector yields a high-quality consensus structure. This approach is computationally efficient at inference time and inherently robust to input variability. When applied to experimental data of the nuclear pore complex (NPC), our method yielded key structural measurements (a radius of 46 nm and an inter-ring distance of 52 nm) that show strong agreement with high-resolution cryo-EM data and other state-of-the-art SMLM analysis methods, validating the effectiveness of this latent-space-centric approach.

The main limitation of our current method is its structure-specificity, as the Pocafoldas network must be retrained for each new macromolecular target. Future work should focus on overcoming this limitation to enhance broad applicability, by potentially developing generalizable architectures, incorporating richer per-localization features like photon counts, and leveraging higher-fidelity training data.

This research successfully demonstrates the viability and potential of a GDL-based pipeline for SMLM structural analysis. By shifting the complex task of particle averaging from coordinate space to a learned, information-rich latent space, we have established an efficient and robust proof-of-concept for analyzing incomplete structural data.

## AUTHOR CONTRIBUTIONS

DMF and PK initiated and conceptualized the research. DMF designed and implemented the methodological framework and computational analyses. PS contributed to the data simulation and statistical analysis. DH and MS provided access to experimental data, laboratory resources, and contributed to the biological validation. PK secured funding and supervised the project. DMF and PK wrote and revised the manuscript, with input from all authors.

## DATA AVAILABILITY

The experimental datasets used in this study, comprising localization data for NUP107 and 3D DNA-origami tetrahedrons, are available via the 4TU.ResearchData repository at https://doi.org/10.4121/13797686. A demonstration dataset containing simulated tetrahedron structures is available in the project repository at https://github.com/dianamindroc/smlm.

## CODE AVAILABILITY

All code for data simulation, network training and validation is freely available under https://github.com/dianamindroc/smlm.

## FUNDING

This work was supported by the Deutsche Forschungsgemeinschaft (DFG, German Research Foundation) – Project number 440773101.

## COMPETING FINANCIAL INTERESTS

No competing interest is declared.

## Supplementary Note 1: Preprocessed data

**Fig. 10.**
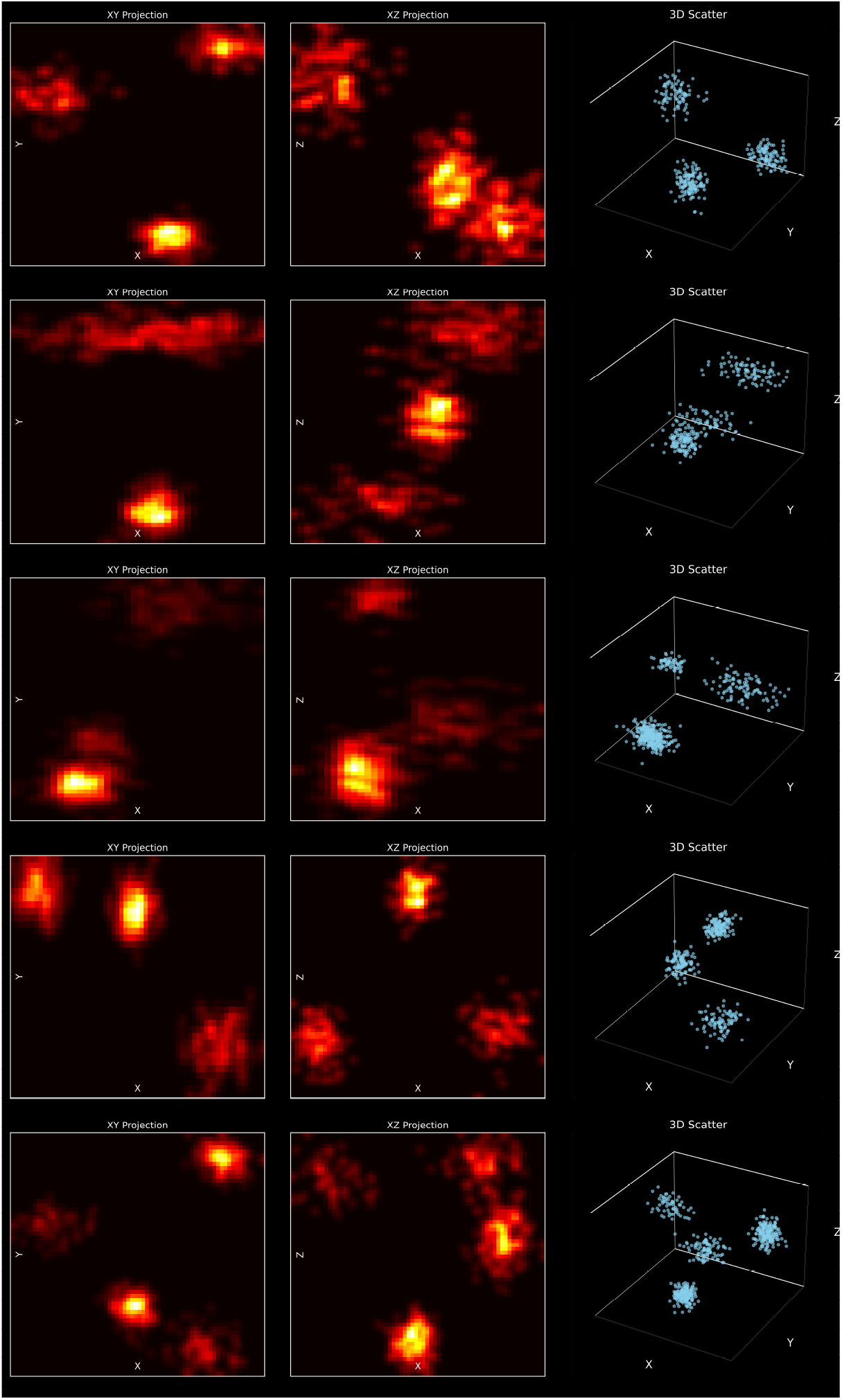
To augment the training dataset and enhance the network’s robustness to experimental variability, the preprocessing pipeline incorporated random rotations and translations. Furthermore, to simulate varying labeling efficiencies and imaging conditions, we applied partial cluster removal and introduced anisotropic noise to the point clouds. These augmented data served as the training input for the network.

## Supplementary Note 2: Attention

**Fig. 11.**
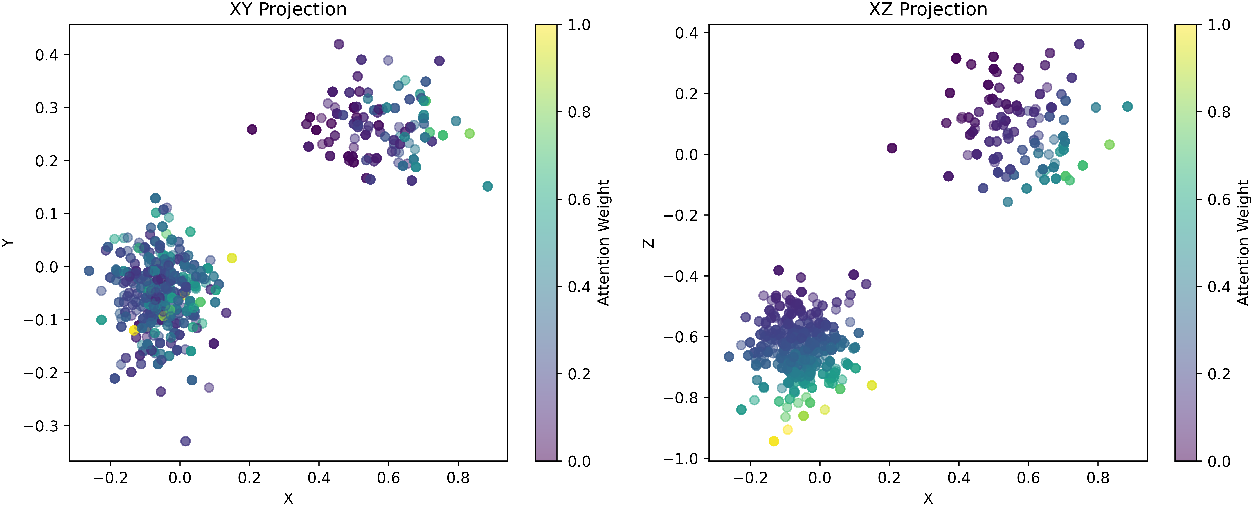
Visualization of the spatial attention mechanism highlights the network’s feature selection strategy. The attention map demonstrates a distinct focus on the peripheral regions of the point clusters, assigning significantly higher weights to points residing on the cluster exterior compared to the interior. This distribution indicates that the network learns to prioritize boundary topology as the most discriminative information for structural inference and reconstruction.

## Supplementary Note 3: Medoid and centroid

**Fig. 12.**
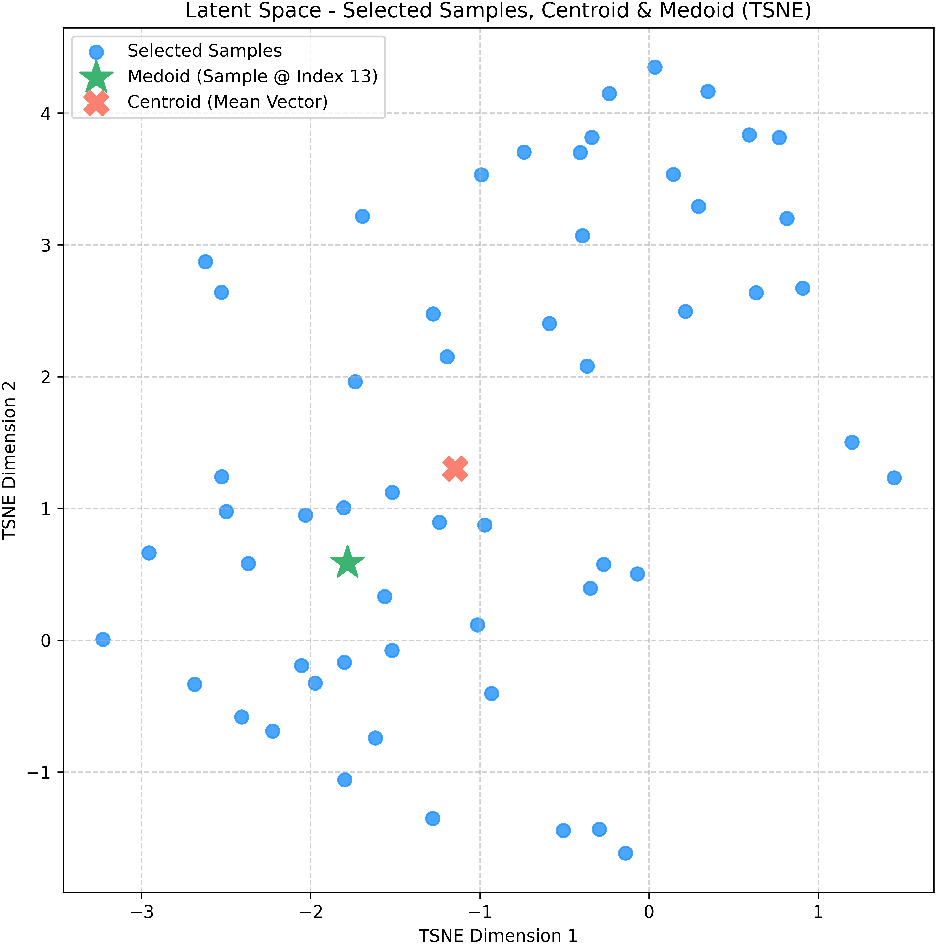
To generate a robust consensus structure, we compute the centroid (red cross), which represents the mathematical mean of all feature vectors in this cluster. However, to ensure the final reconstruction corresponds to a biologically valid conformation within the learned manifold, we identify the medoid (green star)—the specific experimental sample whose feature vector is closest to the centroid in terms of squared Euclidean distance. This medoid serves as the representative ‘average’ particle for subsequent decoding.

## Notes

https://data.4tu.nl/articles/_/13797686

### Competing Interest Statement

The authors have declared no competing interest.

